# Inter-institutional variation in predictive value of the ThyroSeq v2 genomic classifier for cytologically indeterminate thyroid nodules

**DOI:** 10.1101/306126

**Authors:** Andrea R. Marcadis, Pablo Valderrabano, Allen S. Ho, Justin Tepe, Christina E. Swartzwelder, Serena Byrd, Wendy L. Sacks, Brian R. Untch, Ashok R. Shaha, Bin Xu, Oscar Lin, Ronald A. Ghossein, Richard J. Wong, Jennifer L. Marti, Luc G.T. Morris

## Abstract

**Background:** The ThyroSeq v2 next-generation sequencing assay (ThyroSeq) estimates the probability of malignancy in indeterminate thyroid nodules (ITN). Its diagnostic accuracy in different practice settings and patient populations is not well understood.

**Methods:** We analyzed 273 Bethesda III/IV ITN evaluated with ThyroSeq at 4 institutions: 2 comprehensive cancer centers (n=98 and 102), a multicenter healthcare system (n=60), and an academic medical center (n=13). The positive (PPV) and negative predictive values (NPV) of ThyroSeq, and distribution of final pathology were analyzed and compared to values predicted by Bayes Theorem.

**Results:** Across 4 institutions, the PPV was 35% (22-43%), and NPV was 93% (88-100%). Predictive values correlated closely with Bayes Theorem estimates (r^2^=.84), although PPVs were lower than expected. *RAS* mutations were the most frequent molecular alteration. Among 84 *RAS*-mutated nodules, malignancy risk was variable (25%, range 10-37%), and distribution of benign diagnoses differed across institutions (adenoma/hyperplasia 12-85%, NIFTP 5-46%).

**Conclusions:** In a multi-institutional analysis, ThyroSeq PPVs were variable and lower than expected. This is attributable to differences in the prevalence of malignancy, and variability in pathologist interpretations of non-invasive tumors. It is important that clinicians understand ThyroSeq performance in their practice setting when evaluating these results.

## BACKGROUND

Thyroid nodules with indeterminate cytology on fine-needle aspiration (FNA) pose a management dilemma for clinicians, who aim to treat patients with malignant thyroid nodules while avoiding unnecessary medical and surgical therapy in patients with benign disease. Nodules classified as Bethesda III (Atypia of Undetermined Significance/Follicular Lesion of Undetermined Significance) and Bethesda IV (Follicular Neoplasm/Suspicious for Follicular Neoplasm) have estimated malignancy rates of 6-18% and 10-40% respectively; however, these numbers vary markedly between institutions (1, 2, 3). Because there is uncertainty regarding the probability of malignancy in indeterminate thyroid nodules (ITN), molecular tests are more frequently being used as tools to better triage patients to observation or surgery.

ThyroSeq v2 (CBLPath, Rye Brook, NY) is a commonly used molecular test to evaluate malignancy risk in ITN. This assay uses DNA and RNA extracted from FNA cytology material to test for hotspot mutations in 14 genes (*AKT1, BRAF, CTNNB1, GNAS, HRAS, KRAS, NRAS, P1K3CA, PTEN, RET, TP53, TSHR, TERT* promoter, and *E1F1AX*) and 42 gene fusions involving *RET, PPARG, NTRK1, NTRK3, BRAF, IGF2BP3, ALK* and *THADA*, as well as to detect expression changes in selected genes (including overexpression of *MET, PTH, TG*, and *TTF1*, among others) (4, 5, 6).

Several recent single-institution studies have been published evaluating the performance of ThyroSeq v2 (4, 5, 6, 7, 8). These studies, however, are limited by their heterogeneous inclusion criteria and analysis methods as well as their single institution nature, restricting our understanding of this assay’s true diagnostic performance across different practice settings and patient populations. A diagnostic test is expected to have differing performance characteristics (positive and negative predictive values) and accuracy, depending on factors such as the prevalence of disease in the population (9, 10). Therefore, the “real world” performance of ThyroSeq v2 as part of routine clinical care is likely to vary somewhat from institution to institution.

We have previously demonstrated that a gene expression-based classifier for ITN exhibited widely variable performance across different institutions, likely attributable to differences in the prevalence of malignancy in different patient cohorts and variability in pathologist interpretation (10). In order to better understand the performance of ThyroSeq v2 for ITN in routine clinical use across different practice settings and patient populations, we analyzed assay results and matched surgical pathology in patients from our institution and 3 other centers.

## METHODS

We performed a retrospective analysis of 273 Bethesda III or Bethesda IV ITN evaluated in 266 patients with ThyroSeq v2, and subsequently surgically resected, at 4 institutions. These included 98 ITN at a comprehensive cancer center (Memorial Sloan Kettering Cancer Center, New York, NY; MSKCC), 102 from a separate comprehensive cancer center (Moffitt Cancer Center, Tampa, FL; MCC), 13 from an academic medical center (Cedars-Sinai Medical Center, Los Angeles, CA; CSMC), and 60 from at a multi-hospital healthcare system (Mount Sinai Health System, New York, NY; MSHS). The ITN data from MSKCC and CSMC have not been previously published. The data from MCC and MSHS have been previously published but both cohorts were re-analyzed for this study according to the inclusion criteria described below (6, 7). At MSKCC, CSMC, and MSHS, ThyroSeq v2 was ordered selectively on patients, or ordered by outside physicians prior to referral, whereas at MCC it was collected at the time of FNA and performed reflexively for all indeterminate cytology (7). All specimens were collected between 2014-2017 (MSKCC 2014-2017, MCC 2014-2016, CSHS 2015-2017, MSHS 2014-2016). These years reflect the time periods during which each institution conducted their internal reviews of ThyroSeq v2 performance.

All fine needle aspirations (FNA) of the ITN were reviewed by fellowship-trained cytopathologists at the institution where the operation was performed (MSKCC, MCC, CSMC, or MSHS). Postoperatively, surgical pathology and preoperative ultrasound reports were re-reviewed and the biopsied nodule was correlated with findings on surgical pathology by matching the nodule lobe, location within the lobe, and size. Incidental carcinomas separate from the biopsied nodule were considered separately. Seven of the patients included in the analysis had more than 1 ITN evaluated with ThyroSeq. The rates of indeterminate (Bethesda III and IV) category usage at each institution are: MSKCC, 18% of all thyroid cytology specimens; MCC, 26%; CSMC, 15%; MSHS, 18% (11, 12, 13).

ThyroSeq v2 results were considered “ThyroSeq-positive” if alterations with malignancy probability > 30% were reported. ITN with no genetic alterations identified, those exhibiting molecular alterations associated with <30% probability of malignancy, or those with “low frequency” mutations corresponding to allelic fractions ≤5% (for *BRAF, TP53, AKT1, CTNB1, P1K3CA, TERT* promoter, and *RET*) or ≤10% (for *HRAS, KRAS, NRAS, PTEN, TSHR*, and *EIF1AX*) were considered to have a low risk for malignancy and were classified as “ThyroSeq-negative.” The positive (PPV) and negative predictive values (NPV) of ThyroSeq results, and distribution of final pathology were analyzed.

Given the reclassification of non-invasive, encapsulated follicular variant of papillary thyroid cancer as the non-malignant entity “non-invasive follicular thyroid neoplasm with papillary like nuclear features” (NIFTP) in 2017, all non-invasive, encapsulated, follicular variants of papillary thyroid carcinoma were re-reviewed by fellowship-trained head and neck pathologists and re-classified as NIFTP when appropriate (14). Additionally, because of the relatively recent re-classification of NIFTP as a non-malignant entity, PPV and NPV were calculated in duplicate, with NIFTP alternatively considered benign or malignant. Measured PPV and NPV were compared to values predicted by Bayes Theorem (15) based on published prevalence of malignancy of the indeterminate categories at each institution and the weighted test sensitivity/specificity of the Bethesda III and IV categories reported by Nikiforov et al (4, 5, 11, 12, 13). To perform Bayesian analysis with NIFTP considered benign, the published malignancy prevalence for the indeterminate categories at each institution was adjusted by the proportion of nodules that were NIFTP at that institution. Statistical analysis was performed using GraphPad v.7. This study was approved by the Institutional Review Board of Memorial Sloan Kettering Cancer Center.

## RESULTS

Of the 266 patients included, 75% were female (range, 71-79%) with a mean age of 53 years (range, 42-56 years) **(Table 1)**. Of 273 nodules, the mean size was 2.7cm (2.4cm Bethesda III, range 2.1-2.8cm; 3.0cm Bethesda IV, range 2.2-3.5cm), and malignancy rates were 19% for Bethesda III nodules (range, 0-35%) and 46% for Bethesda IV nodules (range, 20-84%). Of 273 nodules, 155 (57%) had ThyroSeq positive results. The proportion of nodules with ThyroSeq positive results was not significantly different between Bethesda III and Bethesda IV groups (59% vs. 54%; p=.37). The ThyroSeq-negative group included 21 patients with low frequency mutations or a quoted malignancy risk < 30% (7 MSKCC, 11 MCC, 0 CSMC, 3 MSHS) **(Table 2)**.

**Table 1.** Demographics of patients included who underwent surgery for Bethesda III or IV indeterminate thyroid nodules with ThyroSeq v2 testing, by institution. B III, Bethesda III; B IV, Bethesda IV; MSKCC, Memorial Sloan-Kettering Cancer Center; MCC, Moffitt Cancer Center; CSMC, Cedars-Sinai Medical Center; MSHS, Mount Sinai Health System.

|  | **MSKCC** |  | **MCC** |  | **CSMC** |  | **MSHS** |  | **Combined** |  |
| --- | --- | --- | --- | --- | --- | --- | --- | --- | --- | --- |
| **Patients (n)** | **97** |  | **97** |  | **13** |  | **59** |  | **266** |  |
| Female, n (%) | 69 (71) |  | 77 (79) |  | 10 (77) |  | 44 (75) |  | 200 (75) |  |
| Age, mean (SD) | 51 (15) |  | 56 (11) |  | 42 (17) |  | 52 (15) |  | 53 (14) |  |
| **Nodules (n)** | **98** |  | **102** |  | **13** |  | **60** |  | **273** |  |
|  | <u>B III</u> | <u>B IV</u> | <u>B III</u> | <u>B IV</u> | <u>B III</u> | <u>B IV</u> | <u>B III</u> | <u>B IV</u> | <u>B III</u> | <u>B IV</u> |
| Nodules, n (%) | **55**<br>**(56)** | **43**<br>**(44)** | **52**<br>**(51)** | **50**<br>**(49)** | **9**<br>**(69)** | **4**<br>**(31)** | **45**<br>**(75)** | **15**<br>**(25)** | **161**<br>**(59)** | **112**<br>**(41)** |
| Nodule size (cm),<br>mean (SD) | 2.1<br>(1.3) | 2.9<br>(1.3) | 2.6<br>(1.6) | 3.0<br>(1.5) | 2.8<br>(1.1) | 2.2<br>(1.0) | 2.5<br>(1.6) | 3.5<br>(2.1) | 2.4<br>(1.5) | 3.0<br>(1.5) |
| ThyroSeq v2<br>positive, n (%) | 41<br>(75) | 33<br>(77) | 16<br>(31) | 17<br>(34) | 8<br>(89) | 3<br>(75) | 30<br>(67) | 7<br>(47) | 95<br>(59) | 60<br>(54) |
| Malignancy rate,<br>n (%) | 19<br>(35) | 36<br>(84) | 5<br>(10) | 10<br>(20) | 0 | 3<br>(75) | 6<br>(13) | 3<br>(20) | 30<br>(19) | 52<br>(46) |

**Table 2.** Low allelic frequency and low malignancy risk mutations on ThyroSeq v2 considered to be “ThyroSeq negative” by institution. MSKCC, Memorial Sloan-Kettering Cancer Center; MCC, Moffitt Cancer Center; CSMC, Cedars-Sinai Medical Center; MSHS, Mount Sinai Health System.

|  | MSKCC<br>(n=7) | MCC<br>(n=11) | CSMC<br>(n=0) | MSHS<br>(n=3) | **Malignant**<br>**n (%)** |
| --- | --- | --- | --- | --- | --- |
| **“Benign” mutations** |  |  |  |  |  |
| *NIS overexpression* (n=1) | 1 |  |  |  | **0** |
| *PTH expression* (n=2) |  | 1 |  | 1 | **0** |
| **Low allelic frequency mutations** |  |  |  |  |  |
| *EIF1AX* (n=2) | 1 | 1 |  |  | **0** |
| *PTEN* (n=1) |  | 1 |  |  | **0** |
| *RAS* (n=11) | 4 | 6 |  | 1 | **2 (18)** |
| *TERT promoter* (n=1) | 1 |  |  |  | **0** |
| *TP53 indeterminate\** (n=1) |  | 1 |  |  | **0** |
| *TSHR* (n=2) |  | 1 |  | 1 | **0** |
| **Malignant, n (%)** | **1 (14)** | **1 (9)** | **0** | **0** | **2 (10)** |

\*Indeterminate due to poor quality sequencing, suspicious for low allelic fraction mutation.

The overall malignancy rate in ThyroSeq-positive nodules (PPV), with NIFTP considered non-malignant, ranged from 22% to 43% across institutions **(Table 3)**. Overall, the PPV of ThyroSeq v2 for malignancy was 35%. The NPV overall was 93%, and ranged from 88% to 100% across institutions. The sensitivity was 87% (range, 73-100%), and the specificity 52% (20%-75%). Using the pre-test probabilities for malignancy of the indeterminate categories at each institution, the predicted PPVs were 79% at MSKCC, 74% at MCC, 83% at CSMC, and 61% at MSHS. The PPVs were lower than predicted, while the NPVs were close to the expected numbers, and there was overall a strong correlation with predicted values (r^2^=.84) (Figure 1). As the sensitivity and specificity of ThyroSeq v2 initially reported by Nikifirov et al were determined before the re-classification of NIFTP as benign, the PPV and NPV of ThyroSeq v2 were re-calculated with NIFTP considered malignant (Table 3, Figure 1). While this raised the PPVs, it also lowered the NPVs, leading to an overall weaker association with expected values (r^2^=.73).

**Table 3.** Predictive Values of ThyroSeq v2 for malignancy, by institution and combined, with NIFTP alternatively considered benign and malignant. MSKCC, Memorial Sloan-Kettering Cancer Center; MCC, Moffitt Cancer Center; CSMC, Cedars-Sinai Medical Center; MSHS, Mount Sinai Health System; PPV, positive predictive value; NPV, negative predictive value.

| Institution | ThyroSeq Status | Malignant | NIFTP | Benign | Diagnostic Performance (NIFTP Benign) | Diagnostic Performance (NIFTP Malignant) |
| --- | --- | --- | --- | --- | --- | --- |
| MSKCC | ThyroSeq-positive (n=74) | 32 (43%) | 30 (41%) | 12 (16%) | PPV-43%<br>NPV-88%<br>Sensitivity-91%<br>Specificity-33% | PPV-84%<br>NPV-79%<br>Sensitivity-93%<br>Specificity-61% |
|  | ThyroSeq-negative (n=24) | 3 (13%) | 2 (8%) | 19 (79%) |  |  |
| MCC | ThyroSeq-positive (n=33) | 11 (33%) | 5 (15%) | 17 (52%) | PPV-33%<br>NPV-94%<br>Sensitivity-73%<br>Specificity-75% | PPV-48%<br>NPV-87%<br>Sensitivity-64%<br>Specificity-78% |
|  | ThyroSeq-negative (n=69) | 4 (6%) | 5 (7%) | 60 (87%) |  |  |
| CSMC | ThyroSeq-positive (n=11) | 3 (27%) | 1 (9%) | 7 (64%) | PPV- 27%<br>NPV-100%<br>Sensitivity-100%<br>Specificity-20% | PPV-36%<br>NPV-100%<br>Sensitivity-100%<br>Specificity-22% |
|  | ThyroSeq-negative (n=2) | 0 (0%) | 0 (0%) | 2 (100%) |  |  |
| MSHS | ThyroSeq-positive (n=37) | 8 (22%) | 2 (5%) | 27 (73%) | PPV-22%<br>NPV-96%<br>Sensitivity-89%<br>Specificity-43% | PPV-27%<br>NPV-91%<br>Sensitivity-83%<br>Specificity-44% |
|  | ThyroSeq-negative (n=23) | 1 (4%) | 1 (4%) | 21 (91%) |  |  |
| Overall | ThyroSeq-positive (n=155) | 54 (35%) | 38 (25%) | 63 (41%) | PPV-35%<br>NPV-93%<br>Sensitivity-87%<br>Specificity-52% | PPV-59%<br>NPV-86%<br>Sensitivity-85%<br>Specificity-62% |
|  | ThyroSeq-negative (n=118) | 8 (7%) | 8 (7%) | 102 (86%) |  |  |

**Figure 1.**
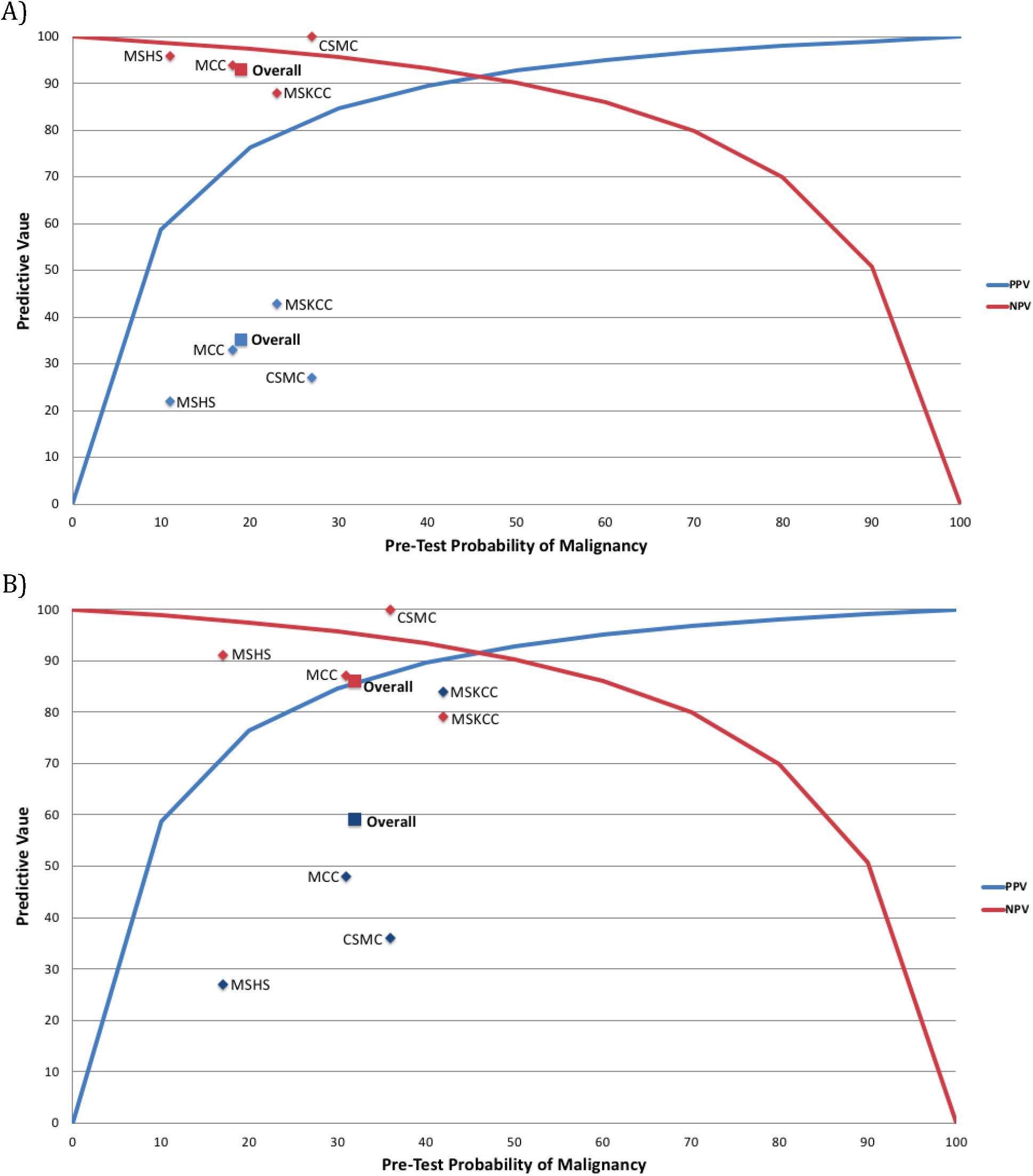
A) Positive and Negative Predictive Values of ThyroSeq v2 compared to values predicted by Bayes Theorem (Pre-test probability; MSKCC 23%, MCC 18%, CSMC 27%, MSHS 11%). B) Alternative values with NIFTP considered malignant (Pre-test probability; MSKCC 42%, MCC 31%, CSMC 36%, MSHS 17%). MSKCC, Memorial Sloan-Kettering Cancer Center; MCC, Moffitt Cancer Center; CSMC, Cedars-Sinai Medical Center; MSHS, Mount Sinai Health System; PPV, positive predictive value; NPV, negative predictive value.

Across all institutions, the most common molecular alterations encountered were mutations of the *RAS* genes, which were found in 62% of all nodules with positive ThyroSeq v2 results **(Table 4)**. There were 96 nodules with *RAS* mutations (64 *NRAS*, 18 *HRAS*, 14 *KRAS*) of which 84 were nodules with isolated *RAS* mutations lacking any other molecular alteration (57 *NRAS*, 14 *HRAS*, 13 *KRAS*). The malignancy rate of all nodules with *RAS* mutations ranged from 9-41% (average 29%). Among *RAS*-mutant tumors, the rate of malignancy was significantly higher when additional molecular alterations were identified (n=12), than when it was found in isolation (58% including only patients with *RAS* + another mutation vs. 25% with isolated *RAS* mutation; p<.05). Other common mutations and their malignancy rates included *BRAFV600E* (5/5; 100%), *BRAF K601E* (1/4; 25%), *E1F1AX* (isolated mutations 2/8; 25%; overall 7/19; 37%), *MET* overexpression (2/7; 29%), *PAX8/PPARG* fusion (5/11; 45%), and *THADA/IGF2BP3* fusion (3/8; 38%).

**Table 4.**
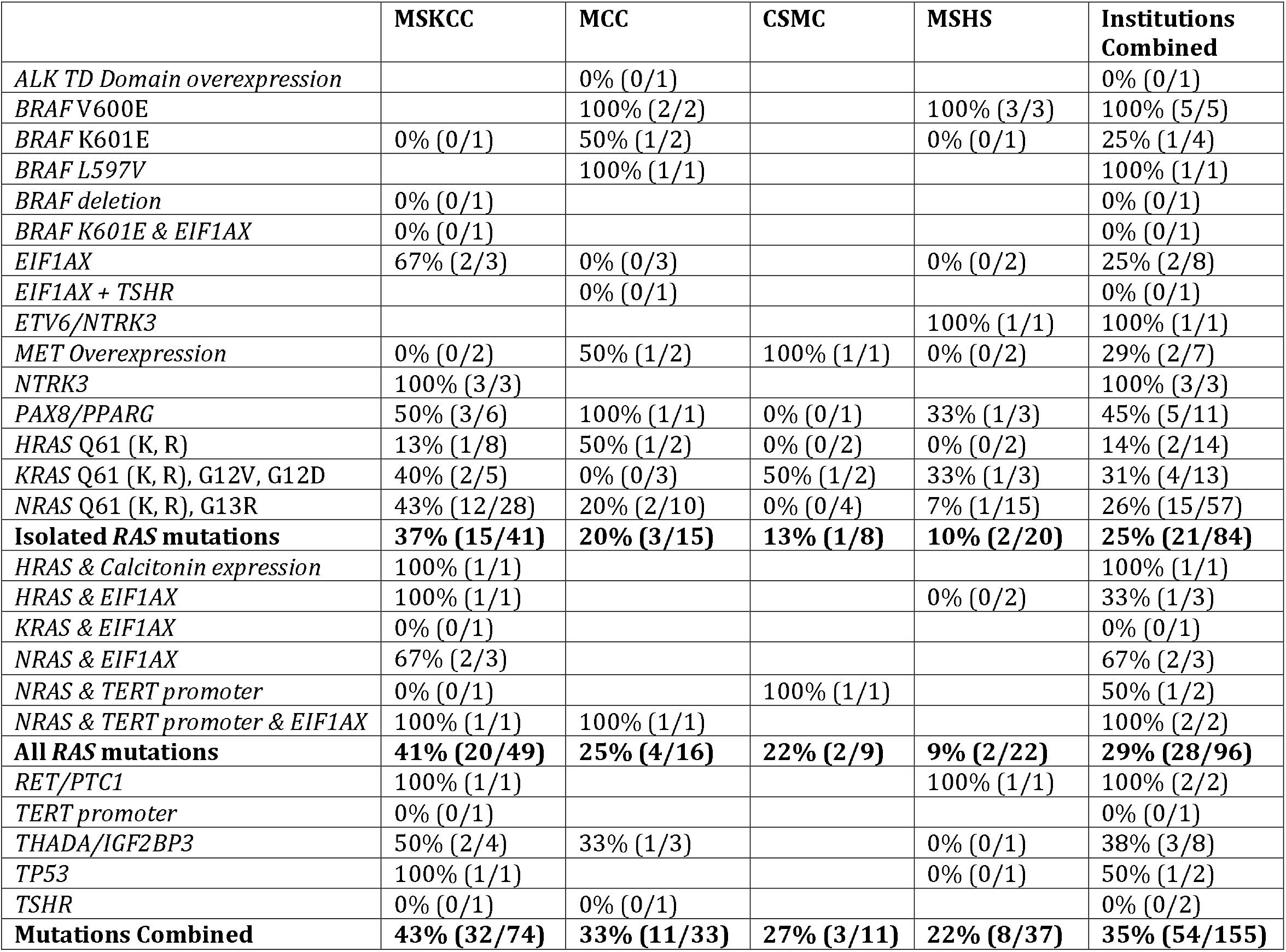
Risk of malignancy with ThyroSeq mutations (resected nodules) by institution MSKCC, Memorial Sloan-Kettering Cancer Center; MCC, Moffitt Cancer Center; CSMC, Cedars-Sinai Medical Center; MSHS, Mount Sinai Health System. NIFTP was considered benign to calculate the rates of malignancy.

The considerable number of *RAS*-mutated nodules in this study allowed for close examination and comparison of their surgical pathology between institutions. At MSKCC, 37% of *RAS*-mutant nodules were malignant (17% classical variant PTC, 10% follicular variant PTC, and 2.4% each of solid variant PTC, mixed classical and follicular variants PTC, follicular thyroid carcinoma, and Hurthle cell carcinoma). Other institutions had lower malignancy rates in *RAS*-mutated nodules: MCC 20%, CSMC 13%, and MSHS 10% (Figure 2). While the *RAS*-mutant nodules at MSKCC had a high rate of NIFTP diagnosis on final pathology (46%), this rate was markedly lower at the other institutions (MCC 7%, CSMC 13%, MSHS 5%), where there were higher rates of other benign pathology, mostly follicular adenoma/nodular hyperplasia (MSKCC 12%, MCC 67%, CSMC 63% MSHS 85%). Across all institutions, *RAS*-mutant nodules were most commonly follicular adenoma/nodular hyperplasia on surgical pathology (44%) followed by NIFTP (26%) and classical variant PTC (11%).

**Figure 2.**
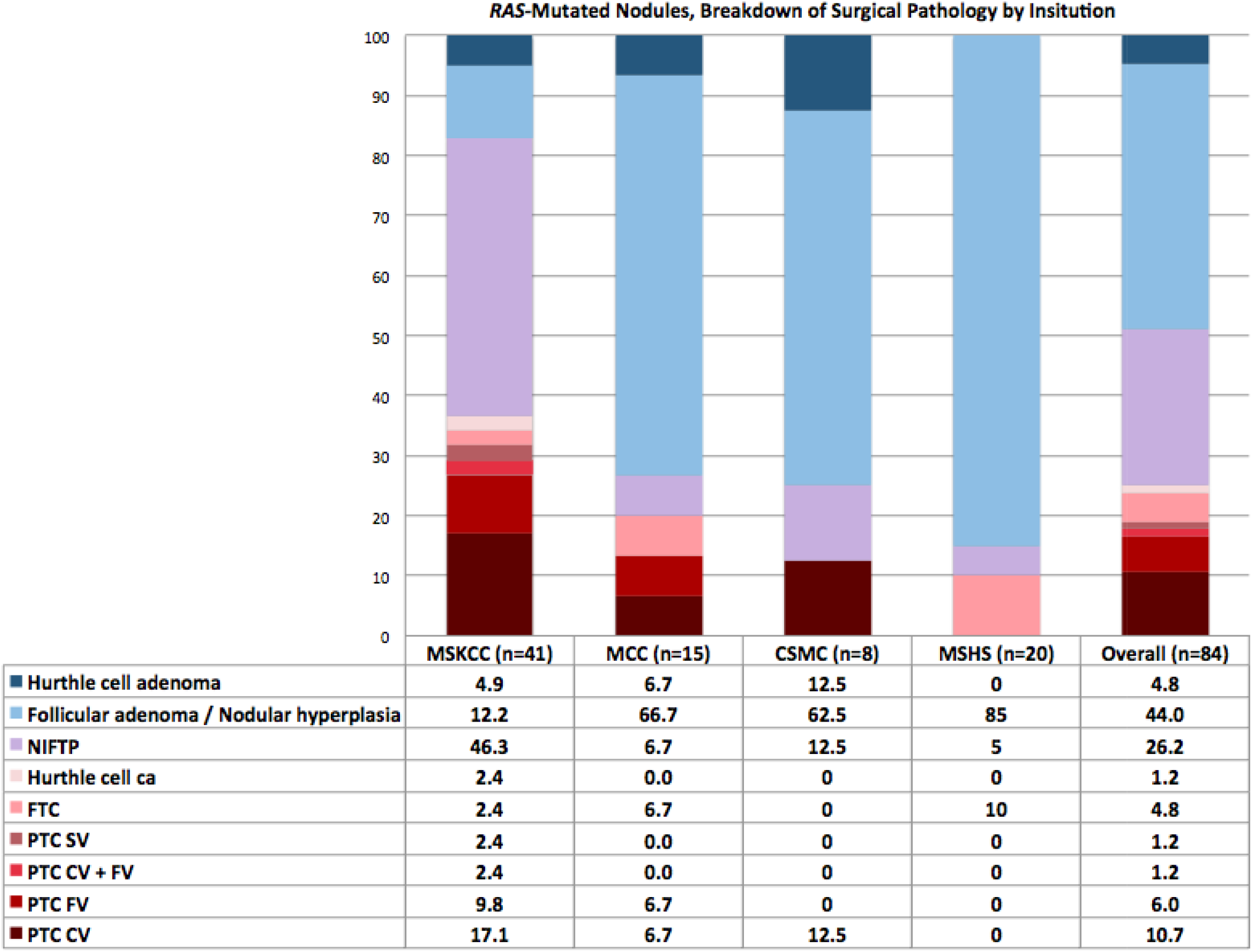
Breakdown of RAS-mutated nodules by surgical pathology and institution, numbers listed as percents. NIFTP, Non-invasive follicular thyroid neoplasm with papillary-like nuclear features; FTC, follicular thyroid carcinoma; PTC, papillary thyroid carcinoma; SV, solid variant; CV, classical variant; FV, follicular variant; MSKCC, Memorial Sloan-Kettering Cancer Center; MCC, Moffitt Cancer Center; CSMC, Cedars-Sinai Medical Center; MSHS, Mount Sinai Health System.

## DISCUSSION

This multi-institutional analysis of ThyroSeq v2 diagnostic performance in 273 ITN reveals marked variation in test performance and the distribution of pathological diagnoses between institutions. While overall test sensitivity was similar to what was originally reported (87% vs. 90%) the specificity was lower than reported (52% vs. 93%) (4, 5). This translated to a wide range of PPVs across institutions (22%-43%), all of which were substantially lower than the initially reported value of 81% (4, 5), and generally lower than the reported probabilities of malignancy on the ThyroSeq report. The NPVs, on the other hand, were less variable (88-100%), with the average NPV of 93% close to the reported value of 96% (4, 5). These findings of a high test NPV and sensitivity, and relatively lower PPV and specificity, reinforce the similar results seen in one prior published series (8).

In addition, we observed wide variation in the prevalence of NIFTP diagnosis (5-46%) across institutions. These results are likely attributable to inter-observer pathologist variability, and underscore the importance of studying the performance of molecular diagnostic assays employed as part of routine clinical care, across different institutions, outside of the more controlled settings afforded by investigational protocols at single institutions.

Even with the relatively low PPV of the test overall, there was a strong correlation with the predictive values estimated by Bayes Theorem, with an r^2^ value of .84. This indicates that the prevalence of malignancy in ITN is an important determinant of PPV and NPV, suggesting that variable disease prevalence is a major cause of variable test performance. When the PPV and NPV were recalculated with NIFTP considered malignant, as it was when the test was designed, there was a weaker correlation with Bayes theorem predictions, with r^2^ of .73. This appears to be attributable to a lowering of the NPV, with a smaller increase in PPV, if NIFTP are considered malignant.

The most commonly detected mutations in this series were the *RAS* mutations, which had a malignancy rate of 25% as an isolated mutation and 29% when patients with all *RAS* mutations were included in the analysis. These values are much lower than the estimated probabilities of malignancy in the ThyroSeq reports - most of these mutations have stated malignancy rates of >70-80%.

There was wide variation in the prevalence of NIFTP in resected ITN across institutions, ranging from 5-46%. At MSKCC, NIFTP was the most common overall and non-malignant entity in *RAS*-mutated nodules. This was substantially higher than the NIFTP rate at the three other institutions, where the most common overall and benign diagnosis was follicular adenoma/nodular hyperplasia. Even between the other three institutions (MCC, CSMC, MSHS), there was significant variation in NIFTP (5-13%) and follicular adenoma/nodular hyperplasia (63-85%) diagnosis rate. Although true differences in histological diagnosis cannot be ruled out due to absence of a centralized pathology review, these results suggest differences in histological interpretation of encapsulated non-invasive follicular lesions as the most likely explanation, which is a well-recognized phenomenon (14, 16, 17).

Even prior to the designation of NIFTP, it had been shown that significant inter- and intra-observer variation existed in distinguishing follicular variant of papillary thyroid carcinoma (FVPTC) from follicular adenoma and follicular thyroid cancer, mostly relying on the identification of nuclear features of papillary carcinoma (pseudo-inclusions, nuclear grooves, ground-glass nuclei) within the lesion, which is not always straightforward. Studies have observed low rates of concordance (39%) among pathologists in the diagnosis of FVPTC (17). Strikingly, the 24 expert thyroid pathologists convened to reclassify encapsulated FVPTC initially agreed on only 1 of 138 cases (14). The addition of NIFTP as a distinct category of neoplasm with follicular and papillary features added an additional layer of complexity to this diagnosis. To attempt to mitigate this inter-pathologist variation, clear histopathologic criteria for NIFTP diagnosis have been established, including a 3 point “nuclear score” and other features; including encapsulation or clear demarcation, follicular growth pattern with <1% papillae, no psammoma bodies, <30% solid/trabecular/insular growth pattern, no vascular or capsular invasion, and no tumor necrosis or high mitotic activity (14). Even with these strict criteria and with pathologist training, there remains marked inter-pathologist variation in the diagnosis of NIFTP versus follicular adenoma (14). Inter-observer variability in NIFTP diagnosis was the major source of inter-institutional variability, even among 4 institutions with high surgical pathology volume for thyroid disease and fellowship-trained specialty pathologists. It is possible that this variation may be wider outside of institutions with high-volume thyroid surgical pathology.

Currently, there is debate as to whether NIFTP is a pre-malignant (18) or benign (14) entity. NIFTP may represent an in situ carcinoma, or hyperplastic proliferations, with evidence of the natural history of untreated NIFTP currently lacking. Unlike other pre-malignant lesions, it is not clear that there are any well-documented cases of NIFTP developing metastatic disease, even at a low rate (14). Interestingly, RAS mutations are observed in follicular adenomas and nodular hyperplasia, similar to the high prevalence of BRAF mutations in benign nevi arising in the skin (19). At present, this issue is unresolved, and it remains unknown whether all NIFTPs need to be surgically treated. If we consider NIFTP benign and not requiring definitive diagnosis or resection, the percentage of RAS-mutated nodules ultimately requiring surgery would be 10-37% (the carcinomas). On the other hand, if we consider NIFTP pre-malignant and requiring surgical resection, the percentage of RAS-mutant cases requiring surgery would vary dramatically across institutions, ranging from 15-83%.

Some of the limitations of this study include its retrospective and multi-institutional design. While efforts were made to exclude incidental microcarcinomas in the surgical pathology specimen, the inadvertent inclusion of these would artificially inflate the PPV. Additionally, intra- and inter-institutional differences in criteria for ordering ThyroSeq v2 exist, as well as in pathologic interpretation of both cytology and surgical pathology. In 3 institutions in this study (MSKCC, CSMC and MSHS), ThyroSeq v2 was sent selectively for nodules in which results would potentially change management. Selective use of a molecular assay may lower the observed PPV and NPV of the test, as nodules that are more likely to be malignant and test positive on ThyroSeq v2 would be triaged directly to surgical resection (bypassing Thyroseq testing), and those more likely to be benign and testing negative on Thyroseq v2 would be observed (bypassing surgery). Though the extent of this effect is unknown, it may contribute to some of the deviation of the predictive values in this study from the values expected on Bayesian analysis. This study sought to audit the performance of this assay in different patient populations, clinician and pathologist practices, and to therefore reflect “real world” use of this assay. The wide inter-institutional variability in performance is likely attributable to these factors.

This analysis of ThyroSeq v2 performance is the largest and most comprehensive since the test’s introduction in 2014, and it helps to reveal inherent differences in test performance between institutions. This variation in test performance is likely attributable to differences in disease prevalence, selection of nodules for molecular testing, and pathologist interpretation of resected nodules. These factors exemplify the importance of distinguishing *efficacy* (results of an intervention or diagnostic test under ideal circumstances) from *effectiveness* (results observed in “real world” clinical practice). It is not uncommon for the performance of a diagnostic test to be reduced in clinical practice, compared to the initial results observed in highly controlled studies. As newer versions of ThyroSeq and other molecular tests are marketed and used more widely, it is important that physicians are proficient in understanding and interpreting these data in the context of PPV and NPV values in their own practice setting, in order to correctly interpret the results and provide optimal patient care.

